# Monitoring Whole-Body Inflammation with Gallium-68 Labeled Polydopamine Iron Oxide Particles via Hybrid Immuno-PET-MRI

**DOI:** 10.1101/2025.02.04.636474

**Authors:** Swannie Pedron, Amaury Guillou, Fabien Fillesoye, Charlène Jacqmarcq, Cécile Perrio, Frank Caruso, Shiyao Li, Yi Ju, Denis Vivien, Maxime Gauberti, Jonathan Vigne, Thomas Bonnard

## Abstract

Monitoring immune reactions via inflammation imaging provides valuable disease prognosis but remains limited by currently available probes and imaging capabilities. In this study, a bimodal imaging probe was developed from microsized polydopamine matrix-based magnetic particles (M3P) functionalized with antibodies that target vascular cell adhesion molecule-1 (VCAM-1) and that are radiolabeled with the radioisotope Gallium-68. This probe combines sensitive inflammation detection via Positron Emission Tomography (immuno-PET) and high-resolution mapping via Magnetic Resonance Imaging (immuno-MRI). The probe represents a powerful disease diagnosis tool to assess the status of systemic inflammation in sepsis as whole-body immuno-PET enables rapid evaluation of the situation, and immuno-MRI provides a more detailed characterization of the identified affected tissues. The elimination profile of the M3P probe is restricted to the mononuclear phagocyte system and does not undergo renal clearance, which is suitable for application in kidney disorders. The efficacy of the hybrid immuno-PET-MRI diagnosis protocol was tested in a murine model of rhabdomyolysis. Kidney inflammation was detected by whole-body immuno-PET, and vascular inflammation patterns could be revealed by high-resolution immuno-MRI. This bimodal PET-MRI probe could significantly improve inflammation monitoring in pathologies characterized by systemic inflammation or lung and kidney dysfunction.

## 1. Introduction

The diagnosis of inflammation is crucial for disease characterization and management yet its assessment remains complex. It is classically easy to evaluate inflammation in external tissues from the observation of clinical signs, or from blood tests via the increase of immune cells or the release of pro-inflammatory cytokines. However, it is more challenging to detect inflammation of internal tissues, but medical imaging techniques such as magnetic resonance imaging (MRI) coupled with gadolinium chelates which highlight hyperemia and inflammatory edema,^[1]^ provide some solutions. Positron emission tomography (PET) also enables visualization of metabolic activity with [^18^F]fluorodesoxyglucose or ligands of translocator protein (TSPO) that reveal microglial activation.^[2]^ However, despite the clear input of these probes for several indications in oncology or neurology, they reveal late signs of the inflammatory process, and the development of a more specific and sensitive approach is warranted.

The blood vessel wall plays a key role in inflammatory mechanisms. The endothelial cells of the intima activate in response to pro-inflammatory stimuli and express adhesion molecules that enable the recruitment of immune cells from the blood to the inflamed tissue.^[3]^ The strategy of immuno-imaging consists of injecting a probe specific to these leukocyte adhesion molecules to reveal their expression.^[4]^ This mechanism sets among the first signs of an immune reaction and such detection provides the earliest diagnosis of inflammation.^[5]^ We previously developed microsized polydopamine matrix-based magnetic particles (M3P) functionalized with antibodies targeting vascular cell adhesion molecule-1 (VCAM-1) that provided high-resolution immuno-imaging on T2* weighted magnetic resonance imaging (MRI) acquisitions.^[6]^ Using microsized particles injected into the bloodstream enables very efficient targeting of inflamed vessel walls and results in a high detection sensitivity for the technique of immuno-imaging. However, MRI modality possesses inherent limitations as the detection of the magnetic probes is not quantitative. Furthermore, a specific tissue needs to be selected for inflammation assessment as whole-body MRI is not compatible with sensitive probe detection.

Positron emission tomography (PET) offers a quantitative and more sensitive probe detection than MRI; up to the picomolar range for PET while limited to the micromolar range for MRI.^[7]^ PET also dispenses fast whole-body scans and is very complementary to MRI.^[8]^ Hybrid PET/MRI is, for this reason, increasingly considered for the diagnosis of inflammation in cardiovascular diseases.^[9,10]^ Gallium-68 is an appealing positron emitter with a high emission energy, a convenient short half-life of 68 minutes and a facilitated accessibility from germanium-68/gallium-68 generators.^[11]^ As other radiometals, it does however require a chelator that forms a kinetically and thermodynamically stable complex with it.^[12]^

Polydopamine is a biomimetic polycathecol self-adherent polymer with multiple features for the synthesis of nanomedicines.^[13]^ It is a suitable material for the generation of the M3P probe as it enables (i) coating and stabilization of the iron oxide nanocrystals, (ii) self-assembly of the iron oxide into microsized polydopamine matrix, (iii) optimized functionalization with antibody via primary amines or thiol groups using Michael addition chemistry or Schiff base reactions,^[14]^ and (iv) fast clearance within the mononuclear phagocyte system.^[6]^ Another useful feature of polydopamine is its capacity to chelate metal ions such as Ga(III) via the formation of hexadentate metal-phenolic complexes.^[15,16]^

In this study, we labelled the M3P magnetic microsized particles with the gallium-68 radioisotope to develop a bimodal probe for hybrid immuno-PET-MRI. We performed immuno-imaging of VCAM-1 expression by using both of these modalities, harnessing their respective advantages to improve disease diagnostic. We tested this novel technology on mice models of sepsis and rhabdomyolysis and showed that we could efficiently monitor lung and kidney inflammation on whole body immuno-PET, and further provide high resolution mapping of vascular inflammation via immuno-MRI.

## 2. Results and discussion

### 2.1. Synthesis of a bimodal PET-MRI probe

#### 2.1.1. Synthesis of microsized matrix-based magnetic particles (M3P)

Iron oxide nanoparticles (ION) were obtained via co-precipitation of Fe(II) and Fe(III) ions at a 2:1 ratio and subsequently self-assembled via dopamine polymerization into microsized matrix-based magnetic particles (M3P) (**Figure 1A**). Particle suspensions presenting a mean hydrodynamic diameter of D_moy_ = 383 ± 39 nm, with an associated polydispersity index of PDI = 0.21 ± 0.03, were obtained (**Figure 1B**). We tested the capacities of the M3P to chelate gallium ions in cold conditions, incubating them at room temperature for ten minutes with an aqueous solution of GaCl_3_ to form [^nat^Ga]Ga-M3P. This did not affect the size distribution, [^nat^Ga]Ga-M3P presented a mean hydrodynamic diameter of D_moy_ = 380 ± 36 nm, with an associated polydispersity index of PDI = 0.19 ± 0.02. Transmission electronic microscopy (TEM) and high-angle annular dark-field (HAADF) images confirmed that the structure of the iron oxide clusters remains unchanged while the X-ray spectroscopy analysis confirmed that gallium could be incorporated into the particles (**Figure 1C,D**). TEM images reveals that the polydopamine iron oxide cluster do not present a smooth surface and an irregular spherical shape, which is commonly observed with such compact assemblies of iron oxide nanocrystals. The elemental analysis also revealed the presence of carbon, nitrogen, oxygen and iron, which is in accordance with the polydopamine and iron oxide composition of the particles. The M3P particles were then functionalized with a rat anti-mouse VCAM-1 antibody (M3P@αVCAM-1) and to the matching isotype control (M3P@IgG) for further experiments. The attachment of the antibodies relies on Schiff base reactions or Michael additions via thiol or primary amine groups from the immunoglobulin to the polydopamine matrix.^[14,17]^ The synthesis protocol of the M3P@αVCAM-1 and M3P@IgG was adapted from previous work, where a comprehensive characterization of similar particles can be found.^[6]^

**Figure 1.**
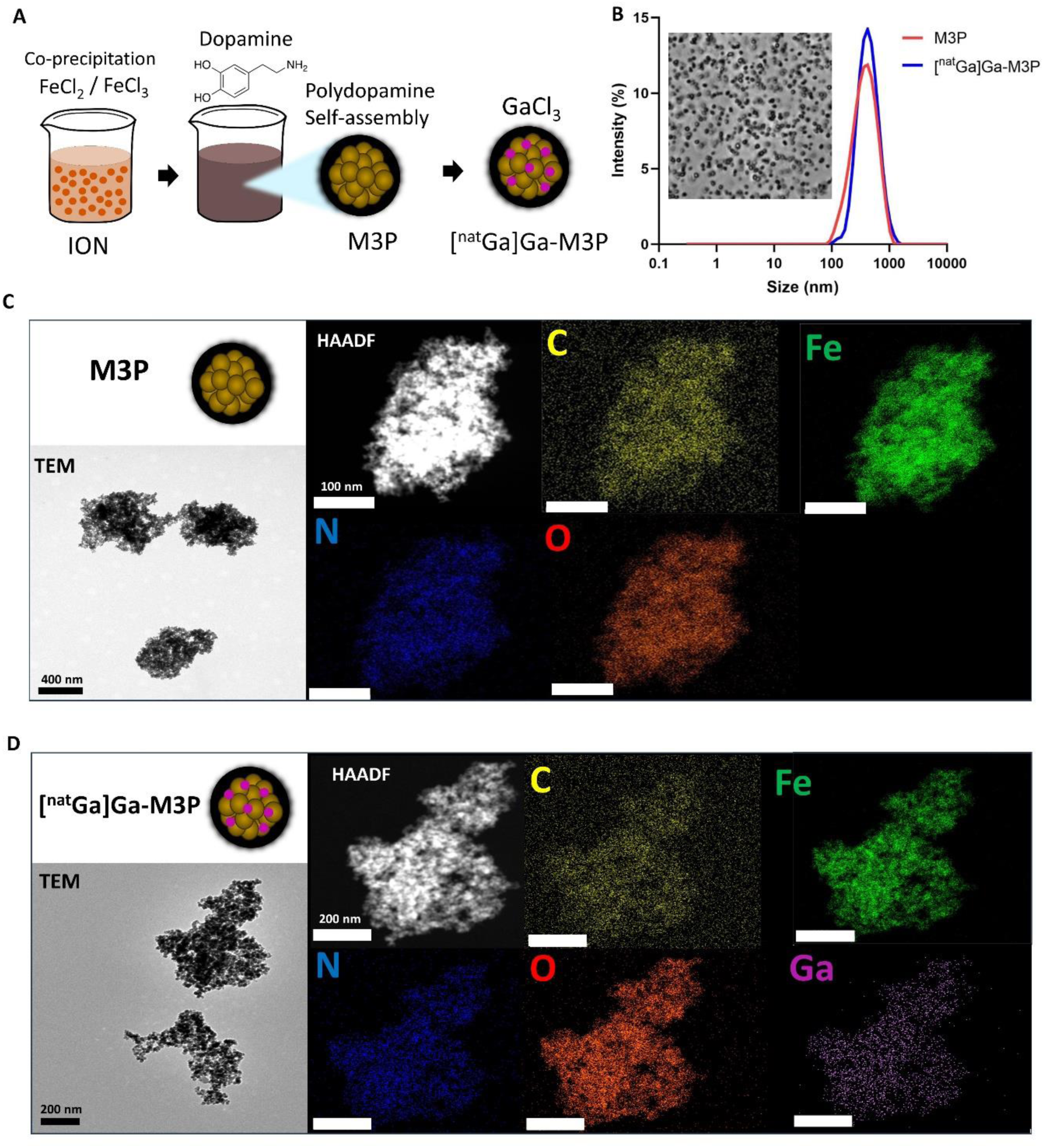
Synthesis of M3P and labelling with gallium ions. A) Synthesis scheme. B) Diffracted Light Scattering measurement of mean diameter distribution by intensity of M3P (D_moy_ = 383 ± 39 nm, PDI = 0.21 ± 0.03) and [^nat^Ga]Ga-M3P (D_moy_ = 380 ± 36 nm, PDI = 0.19 ± 0.02). TEM images, X-ray spectroscopy (EDS) and high-angle annular dark-field (HAADF) images of M3P (C) and M3P labeled with Gallium chloride (D, [^nat^Ga]Ga-M3P).

#### 2.1.2. Radiolabeling particles with Gallium-68 and stability study

Gallium predominantly adopts the +3 oxidation state in an aqueous solution and exhibits properties of a hard Lewis acid. This characteristic facilitates the strong binding of gallium(III) with highly hard Lewis bases, such as carboxylate or phenolate.^[12,18,19]^ The behavior of gallium in chemical reactions is notably influenced by pH, with optimal coordination observed within the pH range between 3 and 5. A more acidic environment may impede reactions by protonating donor atoms, whereas a more basic environment can accelerate hydrolysis, leading to the formation of gallium hydroxide, which precludes its radiolabelling. While most of the ^68^Ga-based radiotracers require the use of a chelating moiety to sequester radio-Ga(III), ION has interestingly shown their ability to strongly chelate ^68^Ga under mild conditions providing radiochemical yield over 90% with strong kinetic inertness.^[20,21]^ Therefore, we studied the ability of our targeted M3P as well as IONs and particles of polydopamine (PDA) to be radiolabelled with the β+ emitter ^68^Ga (**Figure 2**) without the use of a chelating unit. Briefly, M3P@αVCAM-1 or M3P@IgG were resuspended in HEPES buffer (0.3M, 1.5 mL, pH = 4) and (^68^GaCl_3_ ∼500, 1000 MBq) and the mixture was stirred for 5 min at room temperature and the reactions were controlled by radio-ITLC by using citrate buffer (0.1 M, pH = 4.9) as eluant. Using these conditions, all ^68^Ga labeled microparticles ([^68^Ga]Ga-MPs) are found at the baseline of the radio-iTLC (R_f_ = 0 – 0.2), while free ^68^Ga^3+^ present as [^68^Ga]Ga-citrate migrates to the solvent front (R_f_ = 1.0). After magnetic purification to discard uncomplexed ^68^Ga^3+^ and resuspension in mannitol (0.3M), all [^68^Ga]Ga-MPs were obtained in quantitative radiochemical yield (RCYs >98%) and high radiochemical purity (RCPs >99%) (**Figure 2D,E**). It should be noted that the functionalization with antibodies did not affect particle sizes (**Figure 2F**). The incubation in human plasma did not induce any aggregation (**Figure 2G**), confirming results observed in previous studies in stability experiments performed in whole blood.^[6]^ To further validate the use of [^68^Ga]Ga-M3P@mAbs as immuno-PET-MR contrast agents *in vivo* we performed multiple *in vitro* challenge experiments (**Figure 2H**). First the stability towards ligand exchange was carried out using H_4_EDTA as a competitor. Briefly, [^68^Ga]Ga-MPs were incubated with 1000 equivalents of EDTA (0.1 M in chelex-treated water, pH = 7.4) and gently stirred for 15 minutes. Under this condition all the [^68^Ga]Ga-NPs remained stable and is in agreement with other findings.^[20]^ When incubated with either transferrin or Fe(III) as competitor we only found that [^68^Ga]Ga-M3P displayed lower stability compared to [^68^Ga]Ga-ION or [^68^Ga]Ga-PDA and reached 88.9 ± 6.9 and 86.8 ± 6.8%, respectively. Finally, the stability of the radiolabelled NPS in a more biological relevant set up was investigated. As a result after incubation in human serum the different [^68^Ga]Ga-NPs showed full stability thus allowing their further *in vivo* evaluation as multimodal tracers.

**Figure 2.**
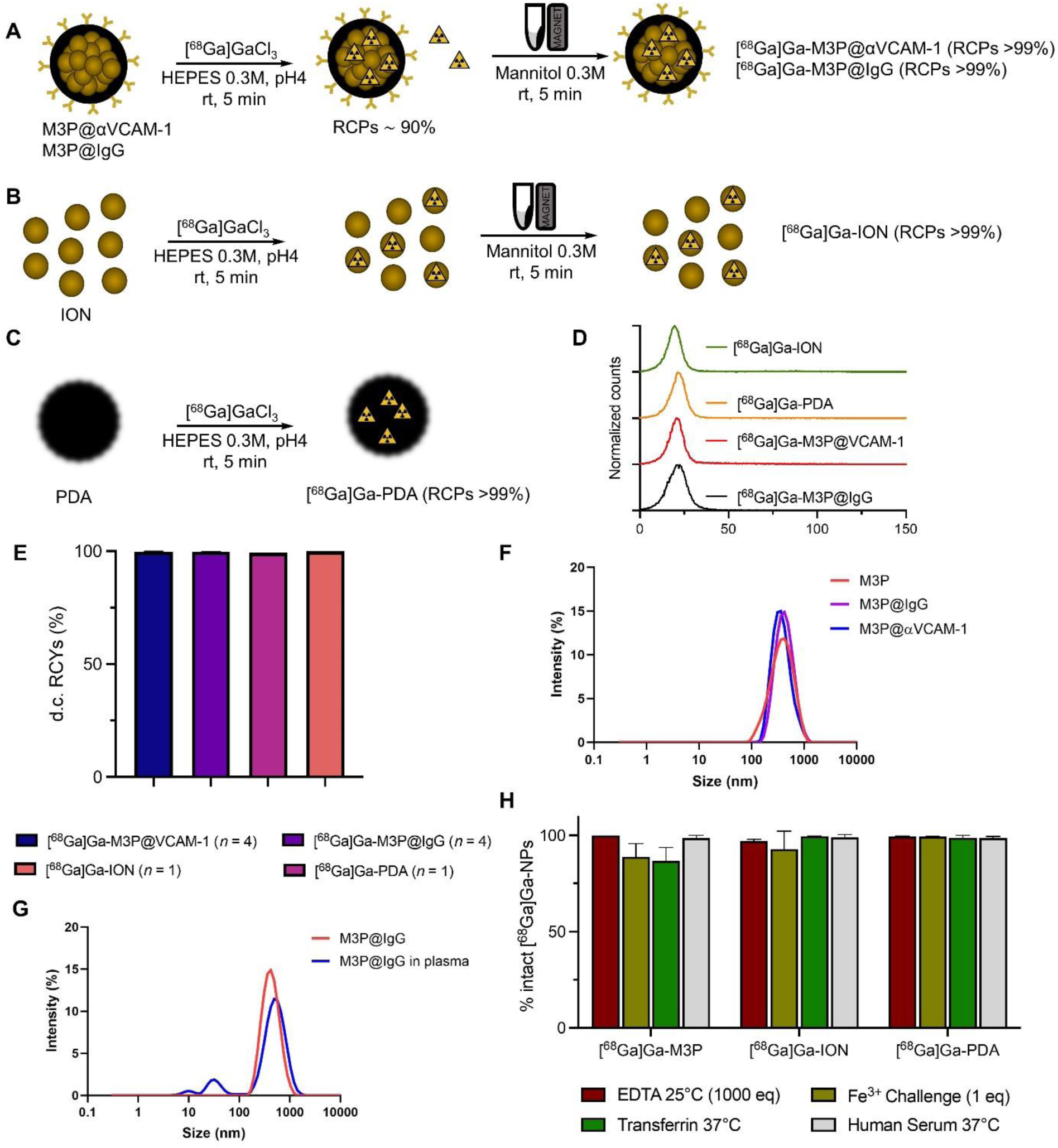
[^68^Ga]Ga-M3P@αVCAM-1, [^68^Ga]Ga-M3P@IgG, ION and PDA radiosynthesis, characterization and *in vitro* stability. A-C) General synthesis scheme for the preparation of [^68^Ga]Ga-NPs. D) radio-iTLC characterization of [^68^Ga]Ga-NPs showing full radiochemical conversion. E) Decay-corrected radiochemical yields of the different [^68^Ga]Ga-NPs. F) Diffracted Light Scattering measurement of mean diameter distribution by intensity of M3P (D_moy_ = 383 ± 39 nm, PDI = 0.21 ± 0.03), M3P@IgG (D_moy_ = 436 ± 47 nm, PDI = 0.27 ± 0.09) and M3P@VCAM-1 (D_moy_ = 407 ± 64 nm, PDI = 0.26 ± 0.10). G) Diffracted Light Scattering measurement of mean diameter distribution by intensity of M3P@IgG in H_2_O or in human plasma. H) Bar chart showing the *in vitro* stability at 15 min post-incubation of the different radiolabelled [^68^Ga]Ga-NPs (*n = 3*).

### 2.2. Whole-body inflammation assessment in sepsis

The inflammation probe was tested in a preclinical positron emission tomography (PET) coupled to a 7 Tesla magnetic resonance imaging (MRI) system (7T PET/MRI, Bruker), in a mouse model of sepsis induced by intraperitoneal injection of lipopolysaccharide (LPS, 5mg/kg) (**Figure 3A**). LPS is an outer membrane component of gram-negative bacteria that induces an innate immune response via stimulation of toll-like receptor 4.^[22]^ This model is characterized by strong systemic inflammation and multiple organ dysfunction, with the lungs, the heart, and the kidneys being the most vulnerable and critical.^[23]^ Sepsis is the principal cause of acute respiratory distress syndrome and commonly induces acute kidney injury.^[24,25]^

**Figure 3.**
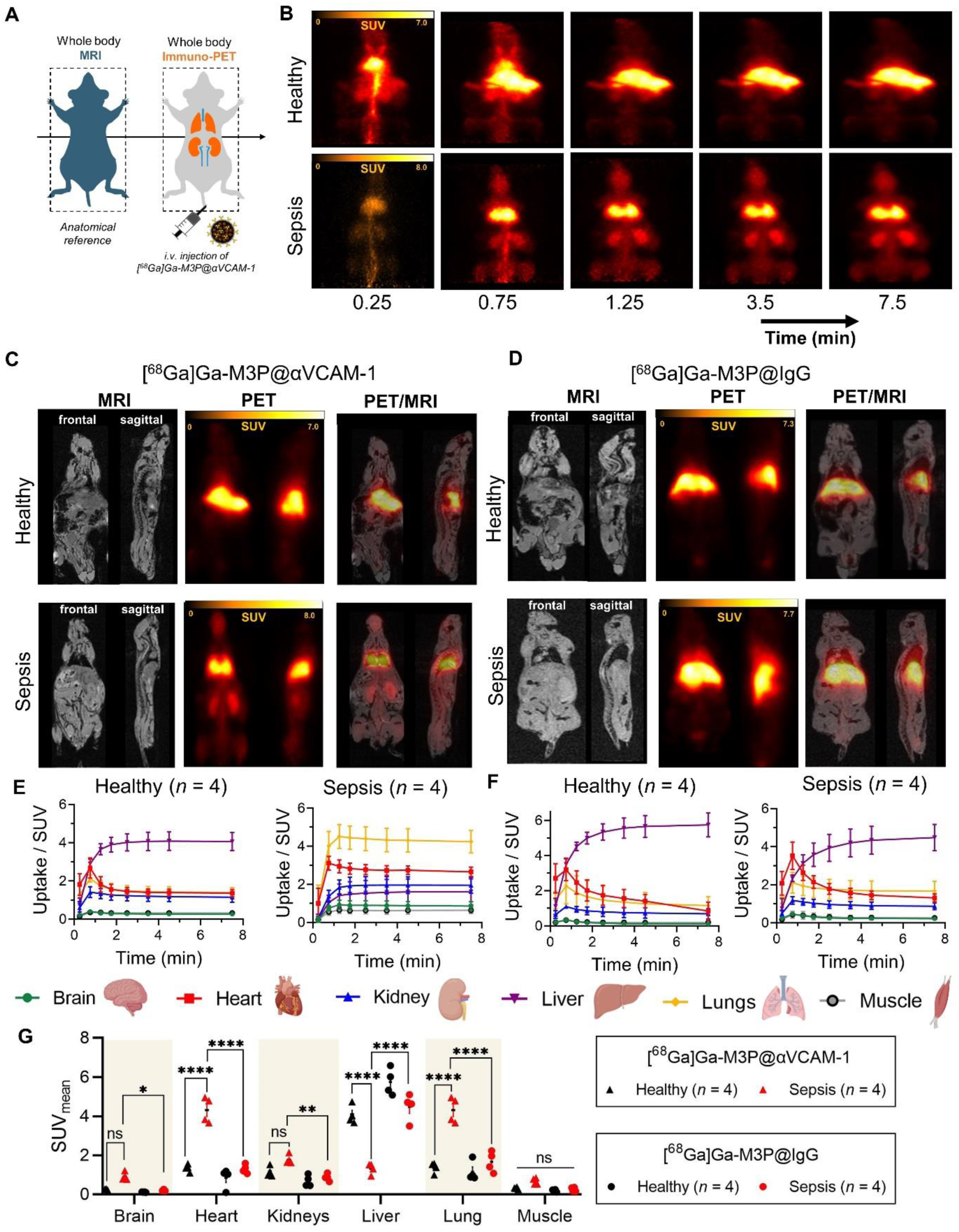
Immuno-PET imaging in a sepsis model. A) Immuno-PET imaging experiment design. B) Dynamic imaging series of immuno-PET acquisition after the injection of [^68^Ga]Ga-M3P@αVCAM-1 in a healthy mouse vs. sepsis mouse. Fused PET-MR acquisition of a healthy control and a sepsis mouse after the injection of [^68^Ga]Ga-M3P@αVCAM-1 (C) vs. [^68^Ga]Ga-M3P@IgG control (D). Corresponding time activity curves in heart, lungs, brain, kidneys, liver, spleen and muscle measured from in vivo PET acquisitions in healthy controls and sepsis mice after the injection of [^68^Ga]Ga-M3P@αVCAM-1 (E) vs. [^68^Ga]Ga-M3P@IgG control (F) (n = 4 / group). G) Standard tracer uptake values quantification from in vivo PET acquisitions (n = 4, two-way Anova analysis with Tukey’s multiple comparison test, *p<0.05, **p<0.01, ****p<0.0001).

The pharmacokinetic profile of the particles radiolabeled with ^68^Ga and functionalized with anti-VCAM-1 antibody ([^68^Ga]Ga-M3P@αVCAM-1) was monitored via whole-body dynamic PET scan (**Figure 3B**) after intravenous injection to healthy and sepsis mice. The first image showing the signal accumulated during the 15 seconds post-injection corresponds to the bolus and thus displays the blood pool. The heart and the main arterial branches such as the abdominal aorta and the carotid arteries are visible for this reason. This is in accordance with the short circulation half-life measured between 50 to 70 seconds for these particles in a previous study.^[6]^In the healthy animals, we noted a fast accumulation in the liver (SUV_mean_ = 4.1 ± 0.5, n = 4), which is in agreement with the extremely short half-life of these particles previously observed due to the fast sequestration by the mononuclear phagocyte system.^[6]^ In the sepsis mice, a strong signal was detected in the lung (SUV_mean_ = 4.2 ± 0.6, n = 4) and in the kidneys (SUV_mean_ = 1.9 ± 0.4, n = 4) indicating that the particles rapidly accumulated in those two specific tissues. To strictly identify the PET signal free from particles in circulation, reconstructions were done with the signal detected from 5 to 10 minutes post-injection and were overlaid with anatomical whole-body MRI scans (**Figure 3C**). As expected, the iron oxide probe is not detected on these whole-body MRI T1-weighted acquisitions as the resolution is too low. When injected with particles functionalized with the matching isotype control antibody ([^68^Ga]Ga-M3P@IgG), no signal uptake other than the liver was noted in both sepsis and healthy mice, (SUV_mean_ = 5.7 ± 0.7, n = 4 and SUV_mean_ = 4.5 ± 0.9, n = 4 respectively) indicating that the signal observed in the sepsis cohort with the [^68^Ga]Ga-M3P@αVCAM-1 is specific for VCAM-1 expression (**Figure 3D**). Giving the short circulation time and the rapid targeting of the probe, the choice of labeling with a relative short lived radioisotope such as gallium-68 is well adapted. PET signal was measured within volume of interest drawn from MRI scan on tissues of interest ,presented as mean standard uptake values (SUV_mean_) and plotted over time post injection (**Figure 3E and 3F**). In healthy mice injected with both particle types and in sepsis mice injected with the non-targeting particles, the signal peaked in the heart at 0.75 minutes post-injection, corresponding to the blood pool bolus signal. In the sepsis mice injected with the VCAM-1 specific particles, the signal remains high after the bolus as the particles bound to the activated endothelium from the inflamed myocardium and thus are not washed away. The same phenomenon is observed in the different tissue as sepsis is characterized by systemic inflammation. The final mean SUV obtained in the different tissues indicated a significant uptake within the brain, the heart, the kidneys, and the lungs of the sepsis mice injected with VCAM-1 targeted particles compared to the sepsis mice injected with the IgG control particles (**Figure 3G**). This suggests inflammation and upregulation of VCAM-1 in those tissues, which is in line with the fact that sepsis is a systemic inflammatory condition. However, it should be noted that the signals in the kidneys and in the brain were not found significantly different from the kidney and brain signals measured in healthy animals injected with the anti-VCAM-1 functionalized particles. We believe that this reveals a limition of our method of analysis when performed at the level of the whole body. Indeed, when comparing various tissues in one single image, the power of the statistical analysis is affected by the inclusion of the other tissues. Besides, when comparing the different signal uptakes isolating every tissues for the analysis, the signal uptake from the brain, the heart, the lung, the kidneys and the muscle of sepsis mice injected with the VCAM-1 targeted particles were significantly higher than in the healthy control injected with the same particles (**Figure S1**). This underlines the interest of a second scan focused on the tissue of interest, as we propose here with the immuno-MRI acquisition, in order to explore more closely the signal differences in various tissues. As a consequence of the targeting of the VCAM-1 particles in the different tissue, the accumulation in the liver is significantly much lower (**** p<0.0001, n=4).

The complete imaging protocol of this study includes whole body T1 weighted MRI acquisition, pre-injection high-resolution T2* kidney MRI, a dynamic PET acquisition of 10 minutes at the beginning of which the probe is injected intravenously, and finally a post-injection high-resolution T2* kidney MRI (**Figure 4A**). We examined more precisely the kidney conditions via immuno-MRI (**Figure 4B**). The iron oxide content of the [^68^Ga]Ga-M3P@αVCAM-1 provided a sensitive contrast in T_2_*-weighted acquisition and enabled high resolution mapping of VCAM-1 expression within the kidneys. We pooled the signal obtained via immuno-PET and immuno-MRI in the kidneys from the different animals for which both were performed and studied the relationship (**Figure 4C**). We measured a strong positive and significant Pearson correlation (r = 0.787, p<0.01) and a linear regression with a good coefficient of determination (R^2^ = 0.619) confirming that both immuno-imaging modalities and the image analysis methods provide reliable and consistent measures of the signal associated to the contrast agent. Signal uptake compare to baseline pre-injection scan was measured and was found significant higher in the kidneys from the sepsis animals injected with [^68^Ga]Ga-M3P@αVCAM-1 (1269.5 ± 486.1 MGU, n = 6) compare to the signal uptake obtained in the kidneys from the healthy animals injected with the same particles (453.6 ± 248.7 MGU, n = 5, p<0.05) or from the sepsis animals injected with the non-targeted control particles [^68^Ga]Ga-M3P@IgG (251.6 ± 257.8, n = 5, p<0.01) (**Figure 4D,E**). VCAM-1 plays a key role in vascular inflammation and is considered as a predictor biomarker of a broad spectrum of cardiovascular diseases.^[26]^ In regards to kidney disorders, recent atlases indicate that VCAM-1 is markedly expressed in response to injury in the epithelial cells from the proximal tubules, in both humans and rodents.^[27,28]^ In the present study, the developed VCAM-1 probe provided a significant signal uptake in the lung, kidneys, heart, and brain. Histological analysis also confirmed that the particles were localized in kidneys from a sepsis animal in the vessel area characterized by VCAM-1 expression, but not in the kidneys from a healthy animal (**Figure S2**). An average of 65 ± 34 particles were found per observed field of view in the kidneys form the sepsis animal versus 13 ± 8 particles per observed field of view in the kidneys from healthy animals (n = 3 / group, ****p<0.0001, **Figure 4F**). In the present histology study, VCAM-1 was only detected on the vessel wall supporting the hypothesis that the VCAM-1 measured with our probe mainly corresponds to the one expressed by the activated endothelial cells. This also aligns with the fact that the probe diameter is too large to cross the endothelium, even when it is damaged and permeable under inflammatory conditions. However, it should still be noted that VCAM-1 may also be expressed by dendritic cells, bone marrow stromal cells, astrocytes, myoblasts, or Sertoli cells for instance. Immuno-labeling of CD68 and Ly6G also revealed a significant increase in the numbers of macrophages and neutrophils respectively in kidneys from sepsis animals compared to healthy control tissue (n = 3 / group, **p<0.01 and ***p<0.001, **Figure 4G**). This marked infiltrate of immune cells indicates that the sepsis kidneys show inflammatory characteristics. We thus confirmed that the PET and MRI signal uptakes noted in the sepsis kidney correspond to an increase VCAM-1 expression and to an inflammatory phenotype.

**Figure 4.**
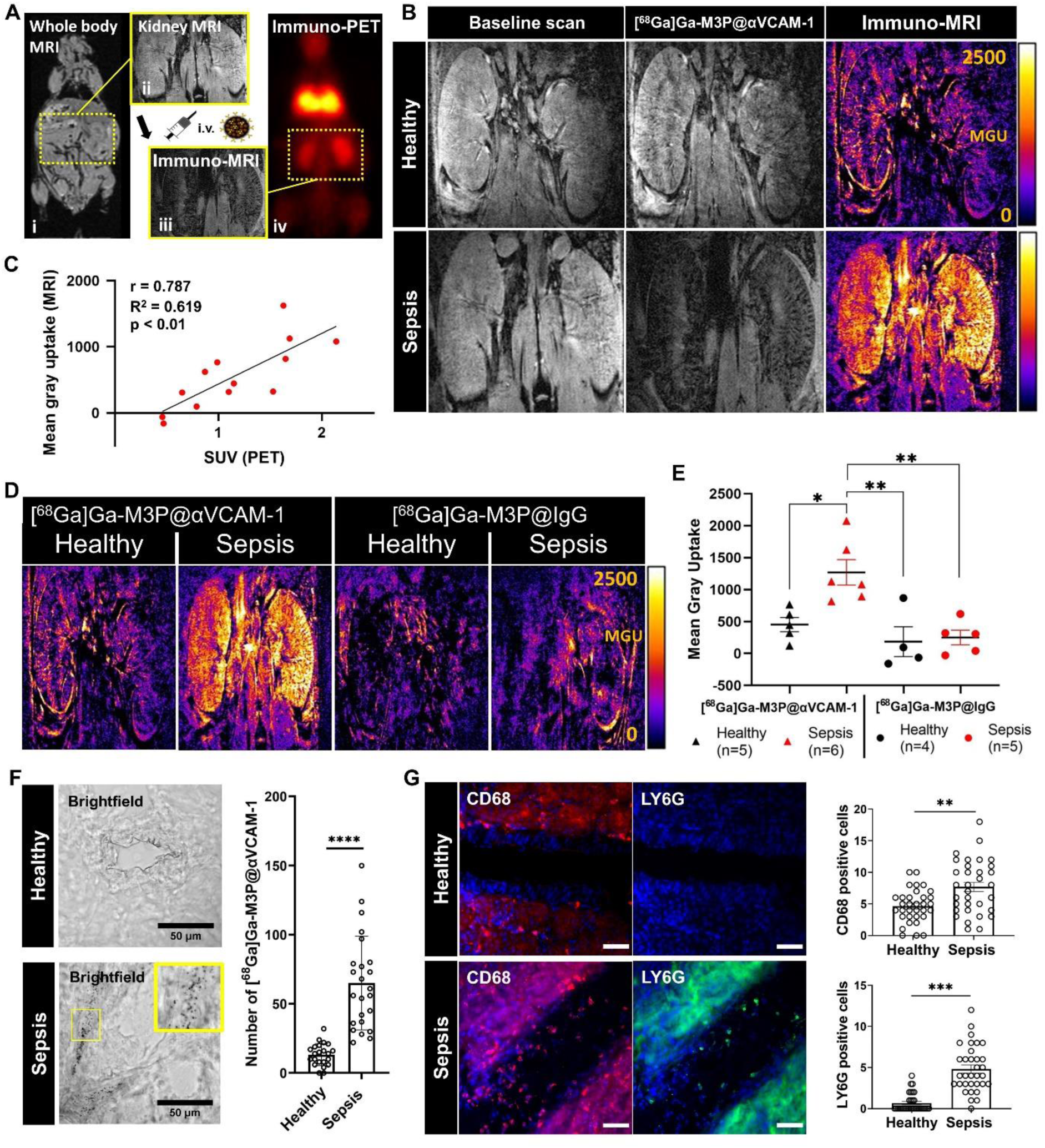
Kidney immuno-MRI in a sepsis model. A) Immuno-PET-MR diagnostic protocol applied to a sepsis mouse; first whole body T1 MRI (i) and baseline kidney T2* MRI (ii) are performed, the [^68^Ga]Ga-M3P@αVCAM-1 or [^68^Ga]Ga-M3P@IgG are then injected intravenously at the begening of the dynamic whole body PET acquisition from which immuno-PET is obtained (iii), and another kidney T2* MRI is finally performed after injection to obtain immuno-MRI (iv). B) Kidney T2* MRI of a healthy and sepsis mouse before (baseline scan) and after the injection of [^68^Ga]Ga-M3P@αVCAM-1. The immuno-MRI image is obtained from image substraction of both MRI to reveal the signal uptake shown with fire look up table. C) Pearson correlation and linear regression between immuno-TEP and immuno-MRI signals obtained in the kidneys. D) Kidnay immuno-MRI of healthy Vs sepsis mice injected with [^68^Ga]Ga-M3P@αVCAM-1 Vs [^68^Ga]Ga-M3P@IgG. E) Kidney immuno-MRI quantifications (n = 4-6, one-way Anova analysis with Tukey’s multiple comparison tests, *p<0.05, **p<0.01. D) Kidney (n = 4, **p<0.01). F) Brightfield observation of kidney slices and quantifications of [^68^Ga]Ga-M3P@αVCAM-1 observed in kidney sections from healthy and sepsis animals and associated quantifications (n = 3 / group, ****p<0.0001). G) Immunostaining of macrophages (CD68, red), neutrophils (LYG6G, green) and DAPI (blue) in kidney sections from healthy and sepsis animals and associated quantifications (n = 3 / group, **p<0.01 and ***p<0.001).

### 2.3. Hybrid immuno-PET-MRI for rhabdomyolysis diagnosis

Although the diagnosis of kidney disorders largely relies on clinical symptoms, assessment of urine samples, glomerular filtration, or biopsy analysis, the implementation of imaging is increasing in nephrology services to refine diagnosis or characterization of kidneys’ structure.^[29]^ In the case of acute kidney injury caused by rhabdomyolysis, a potentially life-threatening condition caused by the breakdown of skeletal muscle cells, there is 1 patient out of 4 that is not clinically diagnosed.^[30]^ Advanced imaging techniques enable observation of muscular atrophy and detection of eventual bleeding or necrosis.^[31]^ Imaging kidney inflammation would provide key information for rhabdomyolysis monitoring and 18-FDG PET/CT is considered for this purpose.^[32]^ Unfortunately, the use of imaging probes in the context of kidney disorder is highly limited as all the current contrast agents available undergo renal clearance, and deciphering the targeted signal from non-specific probe uptake is challenging.^[33]^ This also comes with potential nephrotoxicity side effects that are of particular concern for patients presenting with suspicion of kidney disorders. Thus, using micro-sized biodegradable probes that are exclusively cleared via the macrophages from the mononuclear phagocyte system and do not show any unspecific kidney signal is particularly well adapted for the diagnosis of kidney diseases.

The [^68^Ga]Ga-M3P@αVCAM-1 probe was tested in a rhabdomyolysis model induced by intramuscular injection of glycerol solution. We implemented a serial imaging protocol that enables firstly assessing inflammation at the whole-body level via immuno-PET and secondly imaging more precisely the pattern of vascular inflammation via immuno-MRI focused on the organ of interest (**Figure 5B**). After injection of [^68^Ga]Ga-M3P@IgG, similar signal uptake in the liver was noted in both healthy and rhabdomyolysis animals. This corresponds to non-specific uptake from the macrophages from the mononuclear phagocyte system (**Figure 5C**). However, after the injection of the [^68^Ga]Ga-M3P@αVCAM-1, a specific signal uptake was detected in the kidneys of rhabdomyolysis animals (**Figure 5D**). The quantification as mean standard uptake values indicated a significant signal uptake when compared to the rhabdomyolysis mice injected with [^68^Ga]Ga-M3P@IgG (SUV_mean_ = 1.5 ± 0.3, *n* = 3 Vs SUV_mean_ = 0.3 ± 0.3, n = 3, p<0.01) (**Figure 5E**). High-resolution immuno-MRI acquisitions of kidneys confirmed that VCAM-1 expression could only be detected in the kidneys of rhabdomyolysis mice (**Figure 5F**). The pattern of negative signals on the T_2_* sequences reveals a homogenous vascular inflammation within the radial vessels from the cortical area and the interlobular vessels. The quantification confirmed that a significantly higher signal is obtained in the kidneys of rhabdomyolysis mice injected with [^68^Ga]Ga-M3P@αVCAM-1 (Mean Gray Uptake of 1007 ± 480, n = 3) compared to the signal obtained with [^68^Ga]Ga-M3P@IgG in the kidneys of both healthy and rhabdomyolysis mice (respectively Mean Gray Uptake of -42 ± 129, n = 3, p<0.01 and 177 ± 161, n = 3, p<0.05) (**Figure 5G**). However, VCAM-1 signal difference between the kidneys of the healthy and rhabdomyolysis was less clear and not found significant (p=0.13), which we attribute to VCAM-1 basal expression detected in healthy mice. To date, the few bimodal probes previously developed did not achieve specific imaging of a protein biomarker as we performed here. The various application proposed consist in radiolabeled iron oxide nanoparticles and the imaging approach rely on the passive accumulation of the particles in macrophages or in lymph nodes.^[34,35]^ The present study showcase the first targeted bimodal PET/MRI probe and the sensitive inflammation imaging it enables forshadows marked improvement of disease diagnostic.

**Figure 5.**
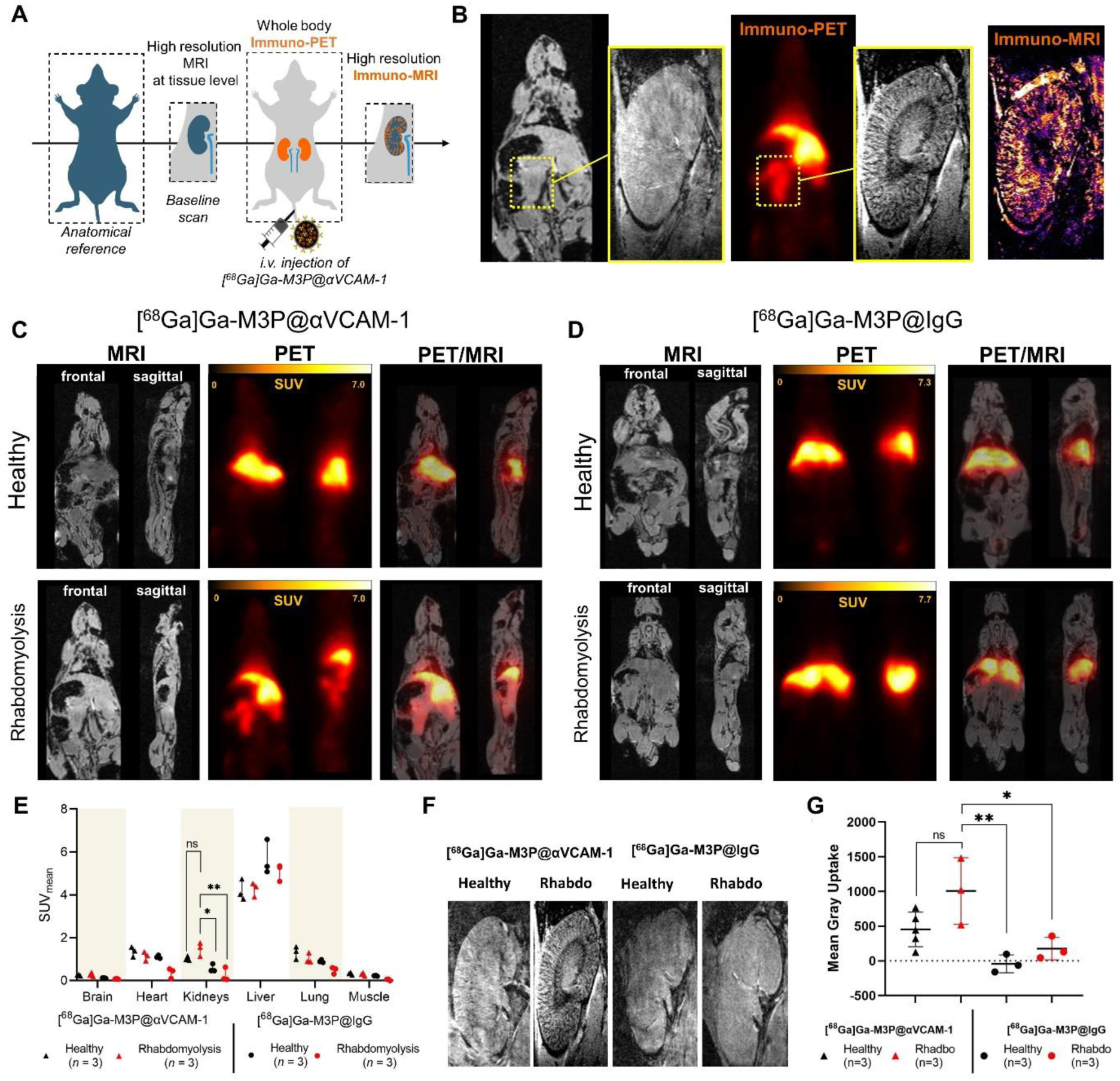
Immuno-PET-MR imaging on rhabdomyolysis model. A) Scheme of the immuno-PET-MR diagnostic protocol. B) Immuno-PET-MR diagnostic protocol applied to a rhabdomyolysis mouse after the i.v. injection of [^68^Ga]Ga-M3P@αVCAM-1. C/D) Fused PET-MR acquisition of a healthy control and rhabdomyolysis mouse after the injection of [^68^Ga]Ga-M3P@αVCAM-1 vs. [^68^Ga]Ga-M3P@IgG control. The healthy control group is the same than presented in Figure 3. E) Standard uptake values ware quantified on PET acquisitions in the different tissues (n = 3 **p<0.01). F) High resolution kidney immuno-MRI of from healthy and rhabdomyolysis mouse after the injection of [^68^Ga]Ga-M3P@αVCAM-1 vs. [^68^Ga]Ga-M3P@IgG control. G) Kidney immuno-MRI quantifications (n = 3,4, ns)

## 3. Conclusion

We reported the synthesis, characterization, and preclinical validation of a novel bimodal PET-MRI inflammation probe. We established a simple and cage-free radiolabelling procedure of the magnetic microsized matrix-based particles (M3P) with Gallium-68 radioisotope, exploiting the metal ion coordination capacities of its polydopamine matrix. Using polydopamine (PDA) based nanomedicine as a metal ion chelator for the synthesis of imaging probe is a popular and efficient strategy. Other groups developed MRI tools from the chelation of gadolinium, manganese or iron ions as the specific electronic relaxation confer signal in T1 or T2 weighted MRI acquisitions ^[36,37]^. In the field of nuclear imaging, the chelation of radionuclides has also previously been performed with polydopamine particles, for instance with Technetium-99m and Iodine-123 for SPECT imaging ^[38]^. In the current study, we introduce the first use of a polydopamine particles with gallium-68 chelation for the development of a PET tracer. The study also differs from the various PDA based imaging tool as we describe a molecular imaging approach using antibody functionalization, applied for the diagnosis of vascular inflammation, distinct from the large majority of PDA nanomedicine developed for cancer applications ^[39]^.

We functionalized the M3P with antibodies directed toward vascular cell adhesion molecule 1 (VCAM-1) or with non-specific immunoglobulin G to obtain non-targeted control probes. VCAM-1 is a leukocyte adhesion molecule with a key role in the regulation of immune surveillance. Monitoring its expression provides an early and sensitive assessment of the inflammation process. In this study, we first showed that the bimodal probe was efficient in revealing lung and kidney inflammation on a whole-body PET scan in a sepsis model. This first step of the diagnosis protocol that we propose, termed immuno-PET, may be a powerful tool in clinical settings to rapidly identify the tissue affected. Sepsis is a complex disease that may affect various tissues with various degrees of severity and such rapid and global assessment of the inflammation status of all tissues may be precious information to guide therapy. We then propose a second step in the diagnosis protocol that consists of looking more precisely at the inflammation pattern in the tissues affected via high-resolution immuno-MRI. We showcased this approach in a rhabdomyolysis model with the imaging of VCAM-1 expression in the kidneys. We demonstrated that the technique could identify vascular inflammation within the radial vessels from the cortical area and the interlobular vessels. Overall, the [^68^Ga]Ga M3P@VCAM-1 carries a high potential to improve the diagnosis of inflammation via PET-MRI. In the context of kidney disorders, this probe has an additional strong asset compared to any other probes currently available as it does not undergo any renal clearance. This results in two significant advantages; the specific signal for inflammation will be much easier to identify accurately in the kidney, and there will be no contraindication due to the risk of nephrotoxicity.

## 4. Experimental Section

### Synthesis of microsized matrix-based magnetic particles (M3P)

The synthesis protocol was adapted from a previous work.^[6]^ Briefly, iron oxide nanoparticles (ION) were obtained via co-precipitation method adding 6.3 mL of 13% ammonia progressively to a 5.7 mL aqueous solution of FeCl_3_.6H_2_O (104.63 mg/mL, Sigma-Aldrich) and FeCl_2_.4H_2_O (142.26 mg/mL, Sigma Aldrich) dissolved in distilled water. The ION were then washed 3 times with distilled water via magnetic separation and resuspended in 40 mL of an aqueous solution of dopamine hydrochloride (2.5 mg/mL, Sigma Aldrich). The ION were then self-assembled into microsized clusters via polymerization of the dopamine induced by the addition of 270 μL of 13% ammonia under continuous stirring for 2 hours using an Ultra-TurraxT-25 disperser at 20,500 rpm. The suspension was then centrifuged at 1000g to remove the large aggregates, the pellet was discarded and the supernatant was washed 3 times with distilled water via magnetic separation and finally resuspended in 8 mL H_2_O. The black suspension obtained corresponds to the microsized matrix-based magnetic particles (M3P). To further functionalized the M3P with antibodies, M3P suspension (1 mL) was incubated with 500 μg of anti-VCAM-1 antibody (M3P@αVCAM-1, A(429), BD Biosciences) or 500 μg match isotype polyclonal IgG (M3P@IgG, Sigma Aldrich) into 5 mL phosphate buffer (10 mM, pH = 8.5) for 24 hours at room temperature under continuous mild agitation. The obtained solution was sonicated for 40 seconds at 20% amplitude and 26 kHz using a UP200ST sonicator tip to break aggregates, washed 3 times with an aqueous mannitol (0.3 M) solution via magnetic separation, and finally resuspended into 5 mL of mannitol (0.3 M). To obtain the micro-sized particles solely composed of polydopamine (PDA), 15 mg of dopamine hydrochloride was added to a 10 mL solution of distilled water with 30% (v/v) ethanol and 0.2 % ammoniac, gently mixed for 6 hours at room temperature.

### hydrodynamic diameter measurement

Dynamic light scattering was used to determine the average hydrodynamic diameter, the polydispersity index and the diameter distribution by intensity of the M3P, [^nat^Ga]Ga-M3P, M3P@IgG and M3P@αVCAM-1 particles with a Nano ZS apparatus (Malvern Instruments, Worcestershire, UK) equipped with a 633 nm laser at a fixed scattering angle of 173°. The temperature of the cell was kept constant at 25°C, and all dilutions were performed in pure water. Stability experiments were performed incubating M3P@IgG in human plasma for 5 minutes before measuring the size. Measurements were performed in triplicate.

### Zeta potential measurement

Zeta potential analyses were realized, after 1/100 dilution in 1 mM NaCl, using a Nano ZS apparatus equipped with DTS 1070 cell. All measurements were performed in triplicate at 25°C, with a dielectric constant of 78.5, a refractive index of 1.33, a viscosity of 0.8872 centipoise, and a cell voltage of 150 V. The zeta potential was calculated from the electrophoretic mobility using the Smoluchowski equation.

### EDS Spectroscopy, TEM, HAADF

Transmission Electron Microscopy (TEM) was performed by a Tecnai F20 instrument (200 kV). Energy-dispersive X-ray Spectroscopy (EDX) mapping was measured by Hitachi SU7000 FE-SEM and Tecnai F20 instrument, respectively. Nanoparticle suspensions were pipetted on and air-dried on plasma-cleaned Formavar carbon-coated copper grids before TEM measurement.

### Radiolabelling procedure

*M3P@αVCAM-1,* M3P@IgG*, ION and PDA particles* were first resuspended in 1.5 mL of 0.3M HEPES buffer (pH = 4). Then, Gallium-68 (500-1000 MBq in 1.1 mL HCl 0.1N) eluted from a ^68^Ge/^68^Ga generator (IRE Elit, Belgium) was directly added to the previous suspension and allowed to incubate for 5 min at RT. Before further use *in vivo*, M3P@αVCAM-1 and M3P@IgG were purified via magnetic separation and resuspended in 2 mL mannitol 0.3M.

### Thin layer chromatography

Glass-fibre iTLC plates impregnated with silica-gel (iTLC-SG, Agilent Technologies) were developed by using an aqueous mobile phase containing citrate buffer (0.1M, pH = 4.9), and were analyzed on an Elysia Raytest Rita Star 2018203 plate reader (Elysia-raytest GmbH, Straubenhardt, Germany). When using aqueous mobile phases containing citrate buffer (0.1 M, pH = 4.9), radiochemical conversion (RCC) was determined by integrating the data obtained by the radio-TLC plate reader and determining both the percentage of radiolabelled product (R_f_ = 0.0) and ‘free’ ^68^Ga (R_f_ = 1.0; present in the analyses as [^68^Ga][Ga(citrate)]). Integration and data analysis were performed by using Gina star TLC software. Appropriate background and decay corrections were applied as necessary. The radiolabeling stability of [^68^Ga]Ga-M3P@αVCAM-1, [^68^Ga]Ga-M3P@IgG, [^68^Ga]Ga-ION and [^68^Ga]Ga-PDA particles were measured in vitro in different conditions.

### Human serum challenge

For each experiment ∼2.5 MBq (50 μL) of the different [^68^Ga]Ga-particles were mixed in 200 μL of human serum. The samples were incubated at 37°C for 2h. The dissociation of the radiolabeled particles was monitored by radio-iTLC (0.1M aqueous citrate at pH = 4.9). Experiments were performed in triplicate.

### Transferrin challenge

For each experiment ∼2.5 MBq (50 μL) of the different [^68^Ga]Ga-particles were mixed with 10 μL of an aqueous Transferrin solution (10 mg/mL). The samples were incubated at 37°C for 2h. The dissociation of the radiolabelled particles was monitored by radio-iTLC (0.1M aqueous citrate at pH = 4.9). Experiments were performed in triplicate.

### H_4_EDTA stability measurements

For each experiment ∼2.5 MBq (50 μL) of the different [^68^Ga]Ga-particles were diluted in Chelex-treated water (50 μL) at pH between 7 and 7.4 followed by the addition of 1000 equivalents (based on iron content) H_4_EDTA (1 mM, pH = 7.4). The samples were incubated at 25°C for 2h. The dissociation of the radiolabelled particles was monitored by radio-iTLC (0.1M aqueous citrate buffer at pH = 4.9). Experiments were performed in triplicate.

### Iron(III) challenge

For each experiment ∼2.5 MBq (50 μL) of the different [^68^Ga]Ga-particles were diluted in Chelex-treated water (50 μL) at pH between 7 and 7.4 followed by the addition of 1 equivalent (based on iron content) FeCl_3_ (100 mg/mL, 10 μL, pH = 7.4). The samples were incubated at 25°C for 2h. The dissociation of the radiolabelled particles was monitored by radio-iTLC (0.1M aqueous citrate buffer at pH = 4.9). Experiments were performed in triplicate.

### Animal models

All experiments were performed on 8- to 10-week-old male Swiss mice (Janvier, France) maintained under specific pathogen–free conditions at the Centre Universitaire de Ressources Biologiques (CURB, Basse-Normandie, France), having free access to food and tap water. Experiments were approved by the local ethical committee of Normandy (CENOMEXA, APAFIS#22318). Sepsis model was induced by a single intraperitoneal injection of LPS (5 mg/kg, Lipopolysaccharides from Escherichia coli O111:B4, Sigma Aldrich) in the lower right quadrant of the abdomen of the animal. The intraperitoneal injection of LPS is followed by a subcutaneous injection of buprenorphine (0.1mg/kg) in order to anticipate any pain or suffering from the animal as soon as possible. The LPS is a pathogen-associated molecular pattern (PAMP), that activates the NF-κB pathway through TLR4 and induces the expression of pro-inflammatory cytokines such as TNFα, IL-6, and IL-1β. Imaging experiments were performed at 24h post-injection of LPS. A systemic inflammation is expected in this model. All sepsis mice were showing signs of disease at 24h; bristly fur, black marks under the eyes and diminished activity. No differences were noted between the different mice from the sepsis group. Rhabdomyolysis model was induced by intramuscular injection of 50% glycerol (7.5 mL/kg, VWR International) in both hind limbs (100 to 150 μL per hind limbs), thereby releasing myoglobin from the muscles in the blood circulation and resulting in acute kidney injury secondary to direct toxicity from myoglobin on the renal tubules. Imaging experiments were performed at 48h post-injection of glycerol. In this model, damages are expected mainly in the kidneys. The mice did not show any external signs of disease and no difference were noted between the different rhabdomyolysis mice.

### TEP-MRI experiments

Experiments were carried out on a small animal 7T PET-MRI system (Bruker, Germany). In total 6 mice groups were studied; healthy controls, sepsis, and rhabdomyolysis injected with [^68^Ga]Ga-M3P@αVCAM-1 and [^68^Ga]Ga-M3P@IgG particles. To reduce the total number of animals required, the same control group of healthy mice was used for comparison with sepsis and rhabdomyolysis. Mice were anesthetized with isoflurane (1.5 to 2.0%) and maintained at 37°C by the integrated heat animal holder, and the breathing rate was monitored during the imaging procedure. A catheter was inserted into the tail vein of mice for intravenous administration of the contrast agent. For anatomical MRI reference, T1_Fisp_3D scans were performed including 3 stitched volumes, allowing to obtain a whole body image of a mice, with the following parameters: 3D, repetition time (TR) 5.5 ms, echo time (TE) 2.6 ms, number of averages (NA) 3, voxel spacing 0.5/0.5/0.5 mm, and a field of view (FOV) 40/40/108 mm. High-resolution T_2_*-weighted images of kidneys for immuno-MRI were acquired with a surface coil (Bruker, Germany), using a 3D FLASH gradient echo imaging (spatial resolution of 78 μm by 156 μm by 300 μm) with TE/TR 8.6 ms/50 ms, and a flip angle of 20°. One baseline scan was performed before the injection of the particles and one after the PET acquisition. List mode PET data were acquired for 10 min, and this was initiated as soon as the formulated [^68^Ga]Ga-M3P@VCAM-1 or [^68^Ga]Ga-M3P@IgG were injected in order to monitor the bolus. The concentration of the probe was maintained between 1 and 4 mg [Fe]/kg and 5 to 20 MBq per animal, giving a molar activity ranging between 2 and 32 Bq/mol. A second similar T_2_*-weighted images of kidneys was finally acquired to detect the magnetic particles accumulated 10 minutes after injection.

### Image analysis

Immuno-PET image data underwent normalization to address discrepancies in PET response, including factors like attenuation, random events, dead-time count losses, positron branching ratio, and physical decay from the time of injection. The Dynamic PET images were reconstructed using an iterative MAP 0.5 algorithm and segmented into 8 different frames (4×30 s; 3×60 s; 1×300 s). To standardize the images in terms of %ID cm-3 (equivalent to %ID/g assuming tissue density as unity), the resulting image data were normalized against the administered activity. Analysis employed PMOD 3.7 software (PMOD, Zurich, Switzerland). Quantification and the creation of time-activity curves (TACs) involved manually drawing 3D volumes-of-interest (VOIs) using fused T1 MR/PET images, with analyses conducted independently by two researchers to minimize variability.To ascertain maximum and average radioactivity accumulation (measured in %ID cm-3 and decay-corrected to the time of injection) across various tissues. Finally, data were transposed into standardized uptake values and were presented as mean values (SUV_mean_) ± SD.

For kidney MRI image data, analysis were performed with ImageJ software (National Institute of Health). Regions of interest were drawn around the kidney’s areas and the mean gray value was measured in the pre-injection baseline scan and in the post-injection scan. The mean gray uptake (MGU) was calculated from the difference between the gray value measured on the baseline and on the post-injection scan. Data were presented as mean values ± SD. The immuno-MRI images were obtained from subtraction of post-injection image to the pre-injection baseline image via the image calculator tool, and the lookup table was changed for fire.

### Immunohistochemistry

Mice were perfused transcardially with saline, the different organs were harvested embedded in a Cryomatrix (Tissue-Tek OCT) and snap frozen via deeping in isopentane chilled in liquid nitrogen. Cryomicrotome-cut transversal sections (10 µm) were collected on poly-lysine slides and stored at -20°C. Sections were co-incubated overnight with rat monoclonal anti-mouse VCAM-1 (1:500; from BD Biosciences), goat anti-collagen-type IV (1:500; Southern Biotech) or with rabbit monoclonal anti-mouse CD68 (1:500; Abcam) and rat monoclonal anti-mouse LY6G (1:200; Biolegend). Primary antibodies were revealed using Fab’2 fragments of Donkey anti-rat and anti-goat to Cy3 and FITC or anti-rabbit and anti-rat to Cy5 and Cy3 (1:800, Jackson ImmunoResearch, West Grove, USA). Washed sections were cover slipped with antifade medium containing DAPI and images were digitally captured using Leica DM6000 epifluorescence microscope coupled CoolSNAP camera, visualized with Leica MM AF 2.2.0 software (Molecular Devices, USA) and further processed using ImageJ 1.49e software (NIH). Cells were considered positive for CD68 or LY6G staining if colocalization with DAPI was observed. The counting of immune cells or M3P on these kidney sections was performed randomly on brightfield images from 3 to 4 sections of 3 different animals, with 3 images per section, to achieve sufficient statistical power. For morphology evaluation, we used a ready-to-use kit for hematoxylin & eosin staining (BioGnost). Briefly, frozen sections were successively rehydrated in deionized water, stained using hematoxylin reagent, rinsed in deionized water, and counterstained with a nuclear bluing reagent. Then, the sections were dehydrated in a gradient bath of ethanol (70, 95, and 100%) and mounted with synthetic resin. Images were digitally captured using a VS120 Virtual Slide Microscope (Olympus) and visualized with QuPath (v0.2.3).

### Statistical analysis

Results are presented as mean values ± SD. Statistical analysis were performed with Graph Pad Prism software (v8.0). Tissues’ SUV_mean_ values measured on immuno-PET acquisitions were compare with two-way Anova followed by Tukey’s multiple comparisons tests. An additional analysis was proposed separating the different tissues and comparing organ per organ with Mann-Withney tests [68Ga]Ga-M3P@αVCAM-1 (Healthy versus Sepsis) and [68Ga]Ga-M3P@IgG (Healthy versus Sepsis). Kidneys’ Mean Gray Uptakes measured on immuno-MRI acquisitions were compared with ordinary one-way Anova followed by Tukey’s multiple comparisons. The numbers of particles functionalized with anti-VCAM-1 antibody versus IgG isotype control in histology sections were compared via a Student’s T test. p < 0.05 was considered significant (two-sided). The correlation between PET and MRI signals in the same animals was studied with a Pearson correlation test.

## Acknowledgments

This work was supported by Agence Nationale de la Recherche “PHySIOMIC” ANR-20-CE19-0031 (to T.B.); “MAD-GUT” ANR-19-CE19-0018 (to S.P. and M.G.); and “Fondation Bettencourt Schueller” for a CCA-Inserm-Bettencourt position (to M.G.). A.G. thanks the European Union’s Horizon 2020 research and innovation programme under the Marie Skłodowska-Curie (grant agreement number 101034329) and the Normandy Region under the WINNINGNormandy program for financial support. C.P. thanks the Normandy Region for further financial support. Y.J. acknowledges support received through the ARC Discovery Early Career Researcher Award scheme (DE230101542). S.P. and A.G. contributed equally to this work. J. V and T. B. contributed equally to this work.

## Competing interests

The authors declare the following competing interests: C.J., D.V., T.B., J.V. and M.G. have filed 2 patents application (WO2023007003A1 and CHIM24197 VIGNE/ACM/AA - EP24306419.3) and for the use of the M3P microparticles as contrast agents. The other authors declare that they have no competing interests.

Received: ((will be filled in by the editorial staff))

Revised: ((will be filled in by the editorial staff))

Published online: ((will be filled in by the editorial staff))

---

Pedron and co-workers report the synthesis of microsized polydopamine matrix-based magnetic particles (M3P) functionalized with antibodies that target vascular cell adhesion molecule-1 (VCAM-1) and radiolabeled with Gallium-68 radioisotope. This bimodal imaging probe enables sensitive detection of lung and kidney inflammation via Positron Emission Tomography (immuno-PET) and high-resolution mapping of endothelial activation via Magnetic Resonance Imaging (immuno-MRI).

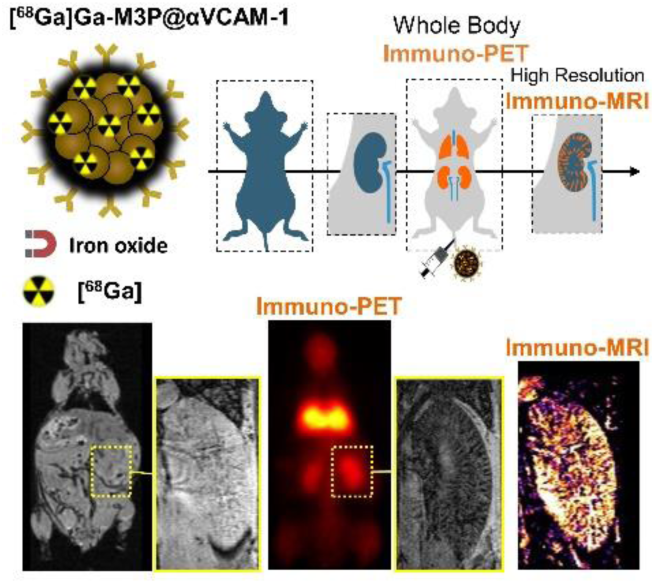

